# An enzymatic assay to measure long-term adherence to pre-exposure prophylaxis and antiretroviral therapy

**DOI:** 10.1101/832410

**Authors:** Ayokunle O. Olanrewaju, Benjamin P. Sullivan, Jane Y. Zhang, Andrew T. Bender, Derin Sevenler, Tiffany J. Lo, Marta Fernandez-Suarez, Paul K. Drain, Jonathan D. Posner

**Affiliations:** Department of Mechanical Engineering, University of Washington, Seattle, USA; Center for Engineering in Medicine, Massachusetts General Hospital, Harvard Medical School, Boston; Department of Materials Science & Engineering, University of Washington, Seattle; Independent Consultant; Department of Epidemiology, University of Washington, Seattle; Department of Global Health, University of Washington, Seattle; Department of Medicine, University of Washington, Seattle; Department of Chemical Engineering, University of Washington, Seattle; Department of Family Medicine, University of Washington, Seattle

## Abstract

Poor adherence to pre-exposure prophylaxis (PrEP) and antiretroviral therapy (ART) can lead to human immunodeficiency virus (HIV) acquisition and emergence of drug resistant infections, respectively. Measurement of antiviral drug levels provides objective adherence information that may help prevent adverse health outcomes. Gold standard drug-level measurement by liquid chromatography/mass spectrometry is centralized, heavily instrumented, and expensive and is thus unsuitable and unavailable for routine use in clinical settings. We developed the REverse TranscrIptase Chain Termination (RESTRICT) assay as a rapid and accessible measurement of drug levels indicative of long-term adherence to PrEP and ART. The assay uses designer single stranded DNA templates and intercalating fluorescent dyes to measure complementary DNA (cDNA) formation by reverse transcriptase in the presence of nucleotide reverse transcriptase inhibitor drugs. We developed a probabilistic model for the RESTRICT assay by calculating the likelihood of incorporation of inhibitors into cDNA as a function of the relative concentrations of inhibitors and nucleotides. We validated the model by carrying out the RESTRICT assay using aqueous solutions of tenofovir diphosphate (TFV-DP), a measure of long-term adherence to PrEP and ART. We used dilution in water as a simple sample preparation strategy to detect TFV-DP spiked into blood. The RESTRICT assay accurately distinguishes TFV-DP drug levels within the clinical range for adherence and has the potential to be a useful test to identify patients with poor adherence to ART and PrEP.

---

For nearly 40 million people living with HIV (PLHIV) and millions more at risk of acquiring HIV,^1^ antiretroviral therapy (ART) and pre-exposure prophylaxis (PrEP) can extend the length and quality of life and prevent HIV infection.^2^ As access to ART and PrEP improves globally, adherence to medication is increasingly becoming a challenge in HIV treatment and prevention.^3–5^ Poor adherence to ART among PLHIV leads to viral rebound, emergence of drug-resistant HIV, and treatment failure.^6^ Poor adherence to PrEP reduces individual and community-level HIV prevention benefits. Roughly 30% of PLHIV receiving ART do not maintain sufficient adherence,^7–9^ and non-adherence rates were higher in several PrEP trials.^10^ Poor adherence arises for several reasons including: presence of barriers to care or medication, medication side effects, psychological problems, and poor provider-patient relationships.^11^ Clinicians, patients, and patient advocates need tools to accurately assess ART and PrEP adherence to effectively implement interventions to improve adherence.^3,12^

There are several approaches for measuring ART and PrEP adherence. Subjective measures of adherence, such as self-reports and surveys,^3,13^ pill counts and tracking of pharmacy refills,^3,4,14^ and wireless pill containers,^15,16^ do not provide proof of actual pill ingestion and have been shown to be inaccurate.^3^ Digital pills with radio frequency transmitters embedded in gel caps provide proof of pill ingestion and information about short and long-term adherence patterns,^17^ but require an individual to wear an RFID receiver that transmits the signal to a cloud-based server, and concerns around cost and privacy could limit acceptance by patients in global settings.^3^

Quantifying concentrations of HIV drugs and their metabolites is an objective approach to measuring drug adherence.^18–21^ Tenofovir disoproxil fumarate (TDF) is used in all PrEP regimens currently recommended by health organizations (e.g. WHO and US Centers for Disease Control) and is also used in ∼58% of all ART regimens.^22^ After ingestion, TDF is hydrolyzed into tenofovir (TFV) and phosphorylated intracellularly by nucleotide kinases into tenofovir diphosphate (TFV-DP), the active form of the drug.^23^ TFV-DP is a nucleotide reverse transcriptase inhibitor (NRTI) that causes DNA chain termination when HIV reverse transcriptase (HIV RT) attempts to form complementary DNA (cDNA) from a RNA template. Tenofovir has a short half-life (17 hours) and a single dose is detectable for up to 4 days in plasma and up to 7 days in urine.^23,24^ Measuring TFV as a marker of adherence is susceptible to the “white coat” effect, where one is unable to distinguish patients who take their medications regularly from those who take their medications just before a doctor’s office visit.^25,26^ On the other hand, TFV-DP has a longer half-life (17 days) and provides cumulative adherence information over a 30-day period.^24,27^ TFV-DP drug levels are associated with health outcomes such as viral suppression in PLHIV,^28^ efficacy of PrEP in HIV-negative people at risk of infection,^29^ engagement in mental health care^20^ among PLHIV, and the risk of viral rebound in PLHIV.^21^

Immunoassays were recently developed to measure TFV in urine and blood.^30–33^ A competitive immunoassay in urine accurately classified 98% of participants in the TARGET study in Thailand who took a dose in the last 24 hours as adherent.^34^ Another competitive immunoassay measuring TFV in urine was tested among PrEP participants at the FIGHT clinic in Philadelphia and distinguished low and high adherence over 48 h and also identified non-adherence that was sustained for more than 7 days before measurement.^33^ All the HIV adherence monitoring immunoassays developed so far have targeted TFV and as such are susceptible to the white coat effect.^25,32^ Developing an immunoassay to detect TFV-DP is challenging since TFV and TFV-DP differ by only two phosphate groups and are both similar to intracellular molecules like adenosine monophosphate and adenosine triphosphate.

TFV-DP drug levels can be measured accurately by liquid chromatography/mass spectrometry (LC/MS).^24,35^ TFV-DP levels in peripheral blood mononuclear cells measured by LC/MS demonstrated that study participants taking ≥ 4 doses/week of PrEP were protected from HIV infection while participants taking ≤ 2 doses/week remained at significant risk of infection.^35^ Castillo-Mancilla *et al* used LC/MS and a pharmacokinetic model to estimate TFV-DP drug levels in red blood cells (RBCs), with median concentrations ranging from 15 – 170 fmol/10^6^ RBCs depending on adherence.^24^ Nevertheless, LC/MS requires significant capital investment, extensive sample preparation, trained personnel, and cold chain storage of reagents and is thus not readily available in clinical settings.^25^ There is considerable interest in developing drug-level assays that can be implemented with minimal capital investment, equipment, and personnel, and that could readily available to physicians.^3,4^

In this paper, we develop an enzymatic assay, termed REverse TranscrIptase Chain Termination (RESTRICT), for ART and PrEP adherence monitoring. The assay takes inspiration from the mechanism of action of TFV-DP on HIV RT and infers drug levels by measuring the length of cDNA generated by RT in the presence of DNA building blocks. Enzyme inhibition assays targeting RT were originally developed in the context of HIV detection,^40–42^ enzyme characterization,^39–42^ drug screening,^43–45^ and drug resistance monitoring.^43,44,46–48^ Although enzyme inhibition assays have been used to evaluate the effectiveness of reverse transcription inhibitors and new drug candidates,^43–45^ to our knowledge such assays have not been applied to measure drug adherence. One reason for this may be that pharmacokinetic data about drug levels corresponding to ART and PrEP adherence were only recently investigated and reported.^24,27^ We formulate a probabilistic model to guide assay design and validate the model with TFV-DP spiked in buffer and blood at clinically relevant concentrations.

## THEORETICAL MODEL

We developed a theoretical model to measure drug concentration using the principles of the RESTRICT assay (Figure 1), The assay requires a nucleic acid template, a primer, free nucleotides (dNTPs), NRTIs (e.g. TFV-DP), intercalating dye, and RT enzyme. RT forms double-stranded DNA (dsDNA) by polymerizing a chain of free nucleotides complementary to a nucleic acid template starting from a region of the template that is hybridized to a primer. At low NRTI concentrations relative to dNTP concentration (scenario A in Figure 1), RT is unlikely to incorporate NRTIs into the cDNA chain and can polymerize the ssDNA into full-length dsDNA strands that bind to many intercalating dye molecules and provide a *high* assay signal. Conversely, at high NRTI concentrations (scenario C in Figure 1), RT is very likely to incorporate TFV-DP into the cDNA chain early, resulting in chain termination and formation of short DNA fragments that bind to fewer intercalating dye molecules and provide a low assay signal. At moderate levels of NRTI (scenario B in Figure 1), the length of the dsDNA product varies and follows a sigmoidal relationship characteristic of enzyme inhibition reactions as shown in Figure 1. In this way, the fluorescence readout from the RESTRICT assay is used to distinguish between low, medium, and high NRTI concentrations.

**Figure 1.**
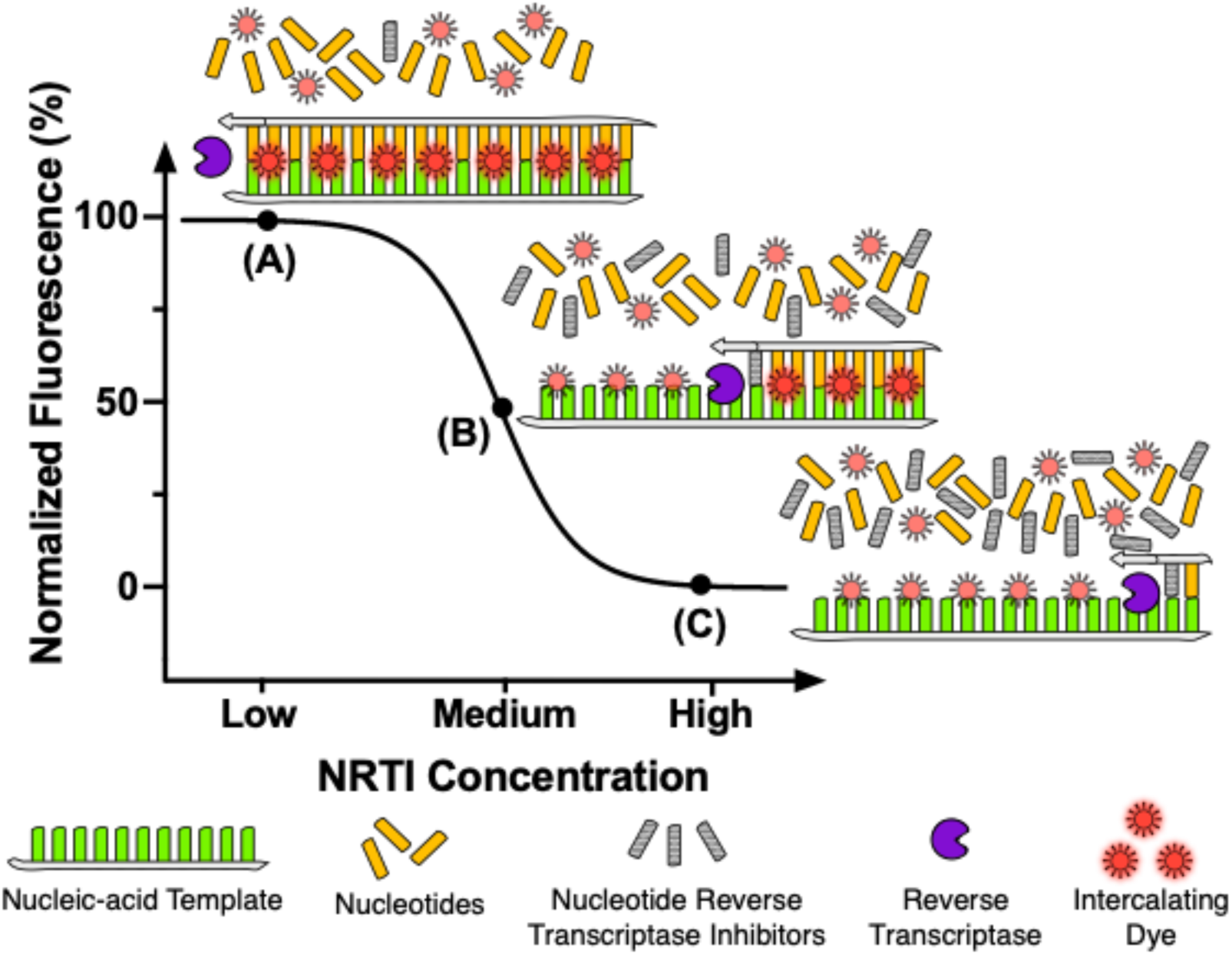
Assay Overview. Schematic illustrating the RESTRICT assay using nucleic-acid template, nucleotides, nucleotide reverse transcriptase inhibitors, reverse transcriptase, and intercalating dye. (A) At low RT inhibitor concentrations relative to nucleotide concentration, reverse transcriptase is most likely to form full-length double-stranded DNA products that provide high fluorescence with intercalating dye, (B) at intermediate RT inhibitor concentrations, fragments of double-stranded DNA that provide intermediate fluorescence are the most likely assay products, while at (C) high RT inhibitor concentration, very little (if any) double-stranded DNA is formed and fluorescence output is due to interactions between the intercalating dye and unpolymerized single-stranded nucleic acid template.

We used DNA templates in the RESTRICT assay because they are less expensive and more stable than RNA templates. It is important to note that although the RESTRICT assay targets the reverse transcriptase (RT) enzyme, our choice to work with DNA rather than RNA templates means that we do not target the reverse transcription function of the enzyme. Instead, the RESTRICT assay targets the DNA polymerization function of RT enzyme. Nevertheless, the polymerization function of the RT enzyme is also inhibited by NRTIs because of the promiscuity of the RT enzyme in incorporating nucleotide analogs during cDNA formation and its poor error correction capabilities.^49^

### Fluorescence from full-length dsDNA products

As illustrated in Figure 1, fluorescence at the end of the RESTRICT assay depends on the interaction between intercalating dye and full-length dsDNA (Figure 1A), dsDNA fragments (Figure 1B), and unpolymerized ssDNA template (Figure 1C). The fluorescence from full-length dsDNA products, *F*_*fp*_, depends on the probability of completion of full-length dsDNA, the length of the DNA template, and the fluorescence properties of the intercalating dye, and can be expressed as,

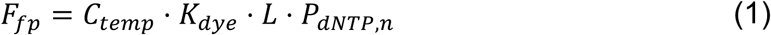

where *P*_dNTP_ is the probability that dNTP is inserted into all successive chain termination sites in the cDNA chain, *L* is the length of a full-length dsDNA product, *K*_dye_ is a constant that represents the fluorescence per double-stranded base pair per unit concentration provided by the intercalating dye, and *C*_*temp*_ is the concentration of DNA template in the assay. In this model, *L* is the number of base-pairs in the entire, full-length dsDNA product, where *n* corresponds to the total number of bases where NRTI could be inserted (i.e. number of bases complementary to the NRTI in the template strand) and is always less than *L*. Both *n* and *L* depend on the exact sequence of the nucleic acid template used in the RESTRICT assay.

We assume that the assay is operating at steady state and that dNTP and NRTIs are not depleted during the assay. In addition, we assume that the probability that dNTP is inserted into each of the *n* available NRTI insertion sites in a DNA template is an independent event. Thus, the probability of formation of full-length dsDNA, *P*_dNTP_, is the probability of a series of *n* successive dNTP incorporation events which can be calculated using the multiplicative rule for probabilities as,

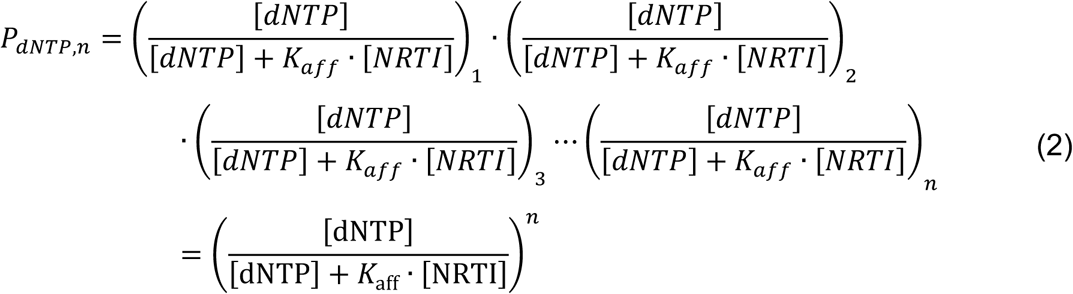

*K*_*aff*_ is the relative affinity of RT for an NRTI compared to its native dNTP substrate, [*NRTI*] is the concentration of NRTI, and [*dNTP*] is the concentration of dNTP present in the assay.

Combining equations (1) and (2), we can express the fluorescence from full-length dsDNA as,

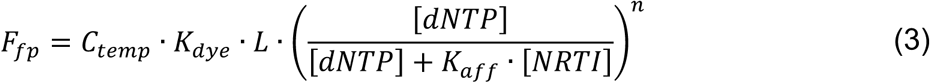

### Fluorescence from fragment dsDNA products

Double stranded DNA fragments also contribute to the total fluorescence at the end of the RESTRICT assay (Figure 1). Similar to full-length DNA, the fluorescence from dsDNA fragments depends on the probability of formation of a fragment and the length of the DNA fragment. At a given dNTP concentration and NRTI concentration, we can calculate the probability of chain termination at any of the *n* possible insertion sites for NRTI in the DNA template. Given that a dsDNA fragment is formed whenever there is an NRTI insertion event, we can calculate the probability of formation of each fragment as,

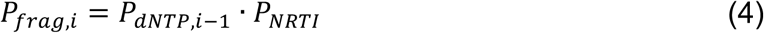

The index *i* counts the bases where it is possible to insert NRTI and the maximum value of *i* is the total number of NRTI insertion bases, or simply *n. P*_*dNTP,i*–1_ is the probability that dNTP is incorporated in the nucleic acid template at all bases preceding base *i* in the template at which NRTI is inserted and can be calculated using equation 3, *P*_*NRTI*_ is the probability of NRTI insertion into the nucleic acid template resulting in chain termination and is simply the probability that NRTI is inserted instead of dNTP and expressed as,

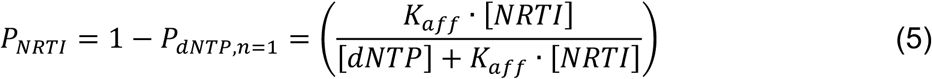

Note that *P*_*dNTP,n=*1_ and *P*_*NRTI*_ are constants for any experimental condition since they represent the probabilities of single insertion events for either dNTP or NRTI, and we assume that we are working in a regime where reagent depletion is not a concern.

We can calculate the probability of formation of fragments of sizes ranging from 1 bp to *n* – 1 bp, where *n* is the total number of available NRTI insertion bases in the DNA template. Given the probabilistic nature of dNTP and NRTI insertion, we expect that there is a distribution of dsDNA fragments sizes at each pair of dNTP and NRTI concentrations. To determine the total fluorescence contribution from dsDNA fragments, we calculate the sum of the fluorescence from all the different dsDNA fragment sizes. Adapting equation (3) and summing the fluorescence from individual dsDNA fragments, the fluorescence from dsDNA fragments can be expressed as follows,

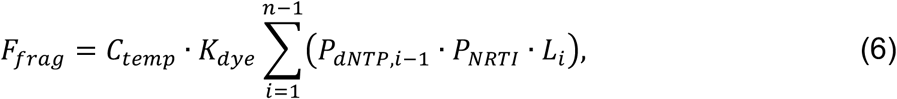

Where *L*(*i*) is the length of the dsDNA fragment when it is terminated at base *i* and can be deduced from the exact sequence of the DNA template.

### Fluorescence from unpolymerized ssDNA template

We also account for the fluorescence contribution of intercalating dye interacting with unpolymerized nucleic acid template. At high NRTI concentrations, very little (if any) dsDNA is formed and most of the fluorescence output comes from interactions between the intercalating dye and the unpolymerized ssDNA template (Figure 1C). Intercalating dyes produce a measurable fluorescence signal when bound to ssDNA fragments. For example, PicoGreen dye used in our experiments, provides 11 times more fluorescence when bound to dsDNA compared to ssDNA.^50,51^ Each dsDNA fragment has a corresponding ssDNA fragment with length equal to the difference between the total length of the nucleic acid template and the dsDNA fragment. Thus, the fluorescence from the ssDNA fragments can be calculated as,

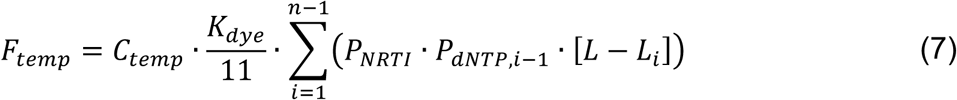

By combining equations (1) – (7) we can calculate the total fluorescence of the RESTRICT assay as,

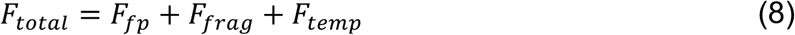

To develop this probabilistic model, we assumed that the assay operates at steady state and that dNTP and NRTI are not limiting reagents in the reaction. For example, the increase in output fluorescence from full-length product as the template concentration increases that is predicted in equation (2) only holds when sufficient dNTP is available to form full-length dsDNA products with all the available nucleic acid template. The overall goal of the theoretical model was not to obtain exact fluorescence values but rather to understand how shape and position of the inhibition curve (Figure 1) changes as assay conditions such as dNTP concentration, TFV-DP concentration, and template concentration change. This allows the probabilistic model to aid in the design of an inhibition assay to quantify TFV-DP within the clinically relevant range of concentrations corresponding to ART and PrEP adherence.

## EXPERIMENTAL SECTION

### RT activity assays

We first determined optimal assay conditions for RT activity in order to minimize assay time and the reagent concentration and cost required. RT enzyme was obtained through the NIH AIDS Reagent Program (Division of AIDS, NIAID, NIH: HIV-1 p66/p51 Reverse Transcriptase Recombinant Protein from Dr. Stuart Le Grice and Dr. Jennifer T. Miller.) Reactions were carried out in an RT assay buffer containing: 60 mM Tris (77-86-1, Sigma), 30 mM KCl (7447-40-7, Sigma), 8 mM MgCl2 (7786-30-3, Sigma), and 10 mM dithiothreitol (20-265, Sigma) buffered to pH 8.0 using HCl (7647-01-0, Acros Organics).

The DNA template consists of a 20 nt primer binding site complementary to the M13 phage DNA primer AGA GTT TGA TCC TGG CTC AG (Catalog, Integrated DNA Technologies, Coralville, IA) followed by TTCA repeats for a total template length of 200 nt and was ordered from Integrated DNA Technologies. For TFV-DP detection, T bases were prioritized in the DNA template because TFV-DP is a deoxyadenosine triphosphate (dATP) analogue and thus will bind to T’s in the DNA template. Although a single-base Poly(T) DNA template would provide maximum indication of TFV-DP inhibition, it was not possible to use a Poly(T) template because HIV RT has a strong preference against Poly(T) DNA templates in polymerization reactions.^52^ Thus, the DNA template was designed using NUPACK software^53^ to preferentially include T bases while also ensuring that the template was free from secondary structures that could lead to unwanted pausing of the RT enzyme.^54^

To characterize RT activity, master mixes consisting of final concentrations of 5 nM DNA template, 5 nM primer, 50 µM deoxynucleotides (dNTPs) (D7295, Sigma), and RT enzyme concentrations of 25, 50, 100, and 200 nM were prepared in black, flat bottom polystyrene 384-well plates with non-binding surfaces (3575, Corning). RT enzyme was added last after which microwell plates were immediately incubated at 37°C in a microplate reader (SpectraMax iD3, Molecular Devices). Assays were stopped by manual addition of 40 µL of PicoGreen intercalating dye (P7581, ThermoFisher Scientific) diluted 1:400 in 1xTE (10128-588, VWR). Reactions were quenched at 16-min intervals up to a total time of 128 minutes. PicoGreen was incubated for 1 min before reading out the assay signal with the microplate reader. Assays were run in triplicate unless otherwise specified.

Data was analyzed using GraphPad Prism 8.1 software (GraphPad Software Inc.). The fluorescence intensity from the RT activity assay as a function of time was fit to an exponential curve. RT activity data as a function of RT concentration was fit to a four-parameter logistic regression curve that follows the familiar symmetrical sigmoidal shape of enzymatic assays. The four-parameter logistic curve fits take the form:

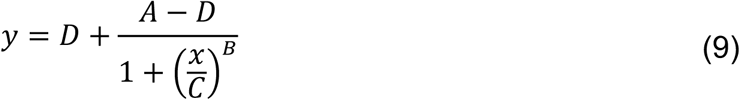

For RT activity assays, *y* represented the fluorescence intensity while *x* represented the enzyme concentration.

### RESTRICT assay in buffer

After determining assay times as well as enzyme and reagent concentrations required for measurable RT activity, we carried out inhibition assays with TFV-DP (166403-66-3, BOC Sciences Inc.). Master mixes for the RESTRICT assay with TFV-DP consisted of 5 µL of DNA template, 5 µL of primer, 20 µL of dNTPs solution, 5 µL of TFV-DP, and 5 µL of HIV-1 RT with reagent concentrations varied to obtain desired final concentrations depending on the experimental conditions tested (see Table S1 in the supplementary information). Serial dilutions of TFV-DP in buffer spanning a concentration range of 1 – 10,000 nM were prepared to generate curves representative of TFV-DP inhibition. Assay characterization experiments were carried out at final dNTP concentrations of 300, 1560, and 6250 nM. Assays were stopped by adding PicoGreen and readout with the plate reader as described earlier.

Fluorescence from RESTRICT assay data was normalized to allow comparison of data points gathered at different dNTP concentrations. Normalization was carried out as,

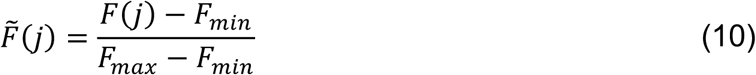

where the subscripts *max* and *min* denote the maximum and minimum measured fluorescence values from each inhibition curve. RESTRICT assay data were fit to four-parameter logistic regression curves that follow the familiar symmetrical sigmoidal shape of enzymatic assays. The 50% inhibition concentration (IC_50_) – the concentration of the drug required to achieve 50% inhibition of its target enzyme *in vitro* was obtained using equation 9 where the parameter *x* is the TFV-DP concentration and the parameter *C* represents the IC_50_. The IC_50_ is often used as a metric to evaluate inhibitor potency and to assess how the position of the inhibition curve shifts (whether lower or higher) when assay conditions (e.g. substrate concentration) are modified.^55^

To compare experimental data and theory, empirical values were derived for the constants *K*_*aff*_ and *K*_*dye*_ (see equation 3). *K*_*aff*_ was chosen by calculating IC_50_ values from experiments and theory at 300, 1560, and 6250 nM of dNTP. *K*_*aff*_ was varied from 0 to 1 in increments of 0.05. We chose a *K*_*aff*_ value that ensured that the experimental and theoretical IC_50_ values differed by < 20% at all dNTP concentrations tested. *K*_*dye*_ was calculated by multiplying the calculated fluorescence from the theory by the ratio of the maximum fluorescence from the experimental data to the maximum fluorescence from the calculated theory.

### RT activity assays in diluted blood

HIV-negative, human whole (BioIVT, Westbury, NY) was diluted in nuclease-free water (3098, Sigma) to lyse RBCs and reduce unwanted inhibition of RT activity by blood components such as immunoglobulins. Blood was mixed with the water by vortexing and incubated for 5 minutes to ensure that the RBCs were lysed. Serial dilutions of blood in water were carried out to make solutions that had final blood concentrations ranging from 2% to 10.0%. 5 µL of diluted whole blood at each final concentration was added to 35 µL of master mix to conduct RT activity assays at 500 nM dNTP. Assays were stopped by adding PicoGreen and read out with the plate reader. Baseline correction was carried out by subtracting the average fluorescence from negative controls (with no RT enzyme and no blood) from the fluorescence obtained from each of the RT activity assays.

### RESTRICT assays in diluted whole blood

We performed RESTRICT assays using blood spiked with TFV-DP. Master mixes for the RESTRICT assay with TFV-DP spiked in 2% blood consisted of final concentrations of 2 nM DNA template, 20 nM primer, 100 nM dNTP, 100 nM of HIV-1 RT, and a range of TFV-DP concentrations. We prepared serial dilutions of TFV-DP in diluted blood to correspond with a concentration range of 5.7 – 11,000 fmol/10^6^ RBCs in whole blood, and thus cover the clinical range for TFV-DP adherence measurement (see Table S2 in the supplementary information). 5 µL of solutions containing TFV-DP spiked in 2% blood were added to 35 µL of master mix so that the final concentration of blood in the RESTRICT assay was 0.25%. Data corresponding to high and low TFV-DP concentrations within the clinical range for adherence measurement were compared using an unpaired t-test in GraphPad Prism.

## RESULTS & DISCUSSION

The RESTRICT assay measures the average length of cDNA synthesized by RT enzyme in the presence of TFV-DP. To demonstrate the utility of this assay for adherence measurement, we first characterized RT activity to ensure that we operated in a regime that provided high assay signal in the absence of TFV-DP. We determined enzyme and reagent concentrations and assay incubation time required for high RT activity. We developed a probabilistic model for the RESTRICT assay and validated this model with experimental data obtained with TFV-DP spiked in aqueous buffer and showed good agreement between experimental data and theory at a wide range of dNTP and TFV-DP concentrations. Finally, we demonstrated that the assay could measure TFV-DP spiked in blood at clinically relevant concentrations.

### RT activity characterization

To characterize RT activity, we first determined the effect of RT concentration and assay time on the rate of cDNA production as measured by the output fluorescence of an RT activity assay. Figure 2A shows a plot of the fluorescence intensity as a function of time in the RT activity assays. At final concentrations of 50, 100, and 200 nM of RT enzyme, the fluorescence intensity increases with time until ∼60 min, when it plateaus. The fluorescence intensity remains flat over time at 25 nM RT. When optimizing enzyme inhibition assays it is desirable to choose an assay time where RT activity provides measurable fluorescence over baseline levels. It is also important to not run the assay for so long that there is a risk of reagent depletion that might skew inhibition measurements. Working at roughly half the time to plateau in an enzyme activity assay typically provides strong measurable signals over background levels and accurate enzyme inhibition measurements.^55^ Thus, we chose a 30 min incubation time for our assay because it provided strong signals over background levels. In addition, the short assay time would also potentially allow for quick (same day) turnaround of adherence measurements to patients and clinicians.

**Figure 2.**
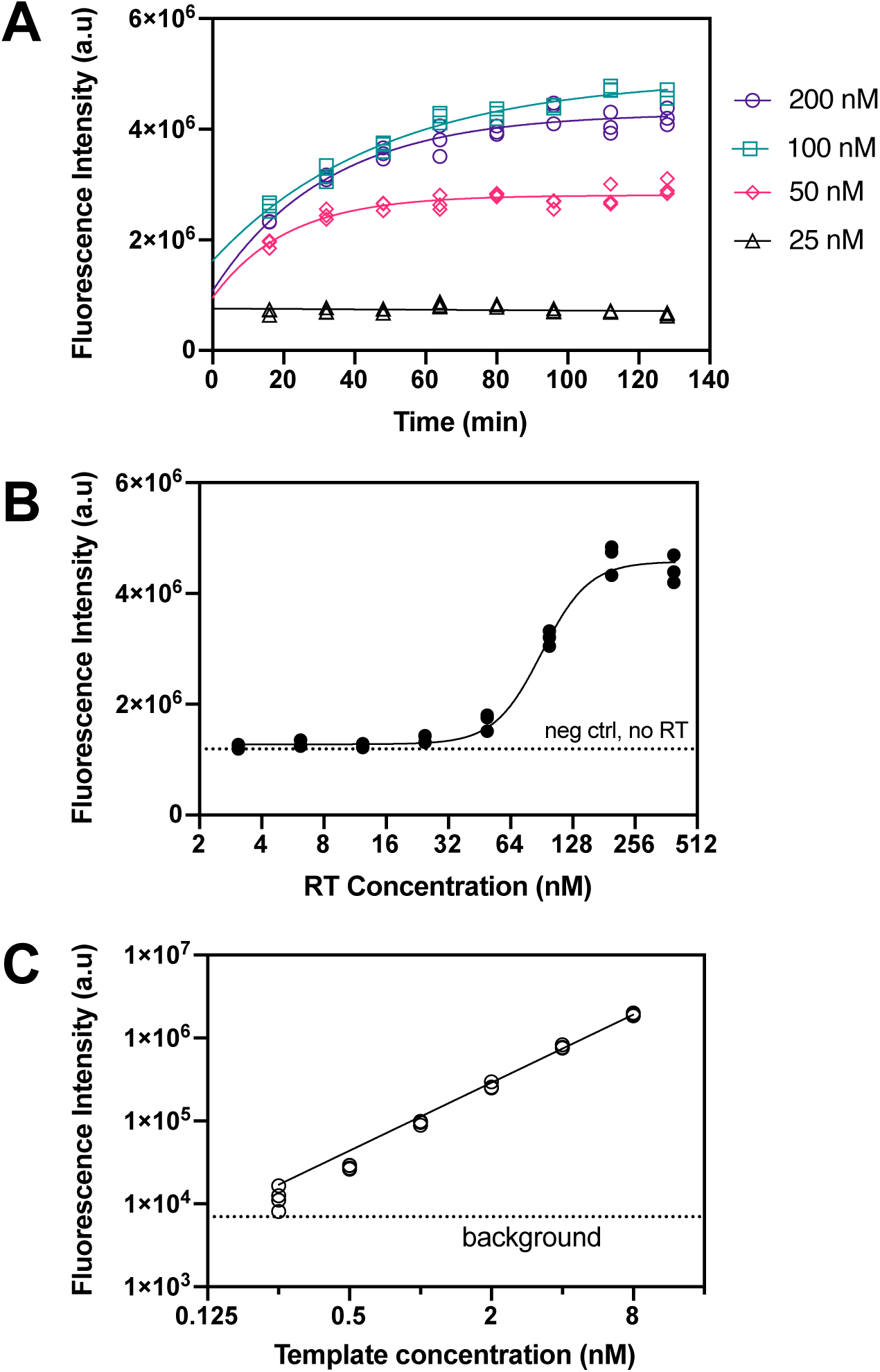
Characterization of RT activity and assay parameters. (A) Measurements of fluorescence intensity over time at different RT concentrations. Lines are an exponential fit of the data. N = 3. (B) The effect of RT concentration on fluorescence intensity at 30 min incubation time. Fluorescence intensity plateaus above 100 nM RT. The line is a four-parameter logistic regression fit of the data. N = 3. (C) The effect of template concentration on fluorescence intensity at 100 nM RT and 30 min incubation time. dNTP concentration was fixed at 200 times the template concentration in each case to ensure that there was sufficient dNTP to form full-length products. The line is a linear fit of the data. N = 4.

To further characterize the effect of RT concentration on assay output signal, we measured the fluorescence intensity at 8 different concentrations of RT enzyme with assay time and reagent concentration held constant. Given that RT is the most expensive reagent used in the inhibition assay, our goal was to determine an RT concentration that provided a strong measurable signal over background levels, while also minimizing the RT concentration required for the assay. Figure 2B plots the endpoint assay fluorescence intensity versus 8 RT concentrations ranging from 3.1 to 400 nM. The fluorescence intensity remains at the same level as the negative control (no RT) until ∼25 nM RT when it begins to increase significantly, and then plateaus above ∼200 nM RT. We chose 100 nM RT as the concentration used in inhibition assays because it provides a large and consistent signal over background levels without using excess RT.

Furthermore, we measured the fluorescence output as a function of DNA template concentration to determine the lowest template concentration and dNTP concentration at which we could measure fluorescence with the plate reader. Figure 2C shows a graph of fluorescence intensity versus DNA template concentration at template concentrations of 0.25, 0.5, 1, 2, 4, and 8 nM. These assays were carried out with corresponding dNTP concentrations of 12.5, 25, 50, 100, 200, and 400 nM, respectively, to keep a consistent 50-to-1 dNTP-to-template ratio across the experiments. There is a linear relationship between template concentration and fluorescence intensity. The lowest template concentration that was detectable above the background signal on our plate reader was 0.25 nM of template. This figure informs how low of a template concentration and dNTP concentration we can use in the RT activity assays in aqueous buffer.

### RESTRICT assays in buffer

We performed assays with TFV-DP spiked in aqueous buffer. Serial dilutions of TFV-DP in buffer spanning a concentration range of 1 – 10,000 nM were prepared to obtain inhibition curves. This concentration range covers two orders of magnitude above and below the clinically-relevant concentration range for TFV-DP adherence described in pharmacokinetic studies.^24,27^ Using the model and the experiments we determined *K*_*dye*_ = 2.03 × 10^12^ as the fluorescence per double-stranded base pair per unit concentration provided by the intercalating dye and *K*_*aff*_ = 0.50 as the relative affinity of RT for TFV-DP compared to its native dNTP substrate.

Figure 3A shows experimental data and theory of the fluorescence from the RESTRICT assay. The data shows the typical sigmoidal-shaped curves representative of enzyme inhibition assays as a function of TFV-DP concentration. In both the experimental data and the theory in Figure 3A, the fluorescence intensity decreases with dNTP concentration. This decrease occurs because we keep the ratio of dNTP and DNA template concentrations constant, to prevent dNTP depletion. Equations 1, 6, and 7 show that the fluorescence from full-product DNA, fragment DNA, and unpolymerized DNA template are directly proportional to the DNA template concentration.

**Figure 3.**
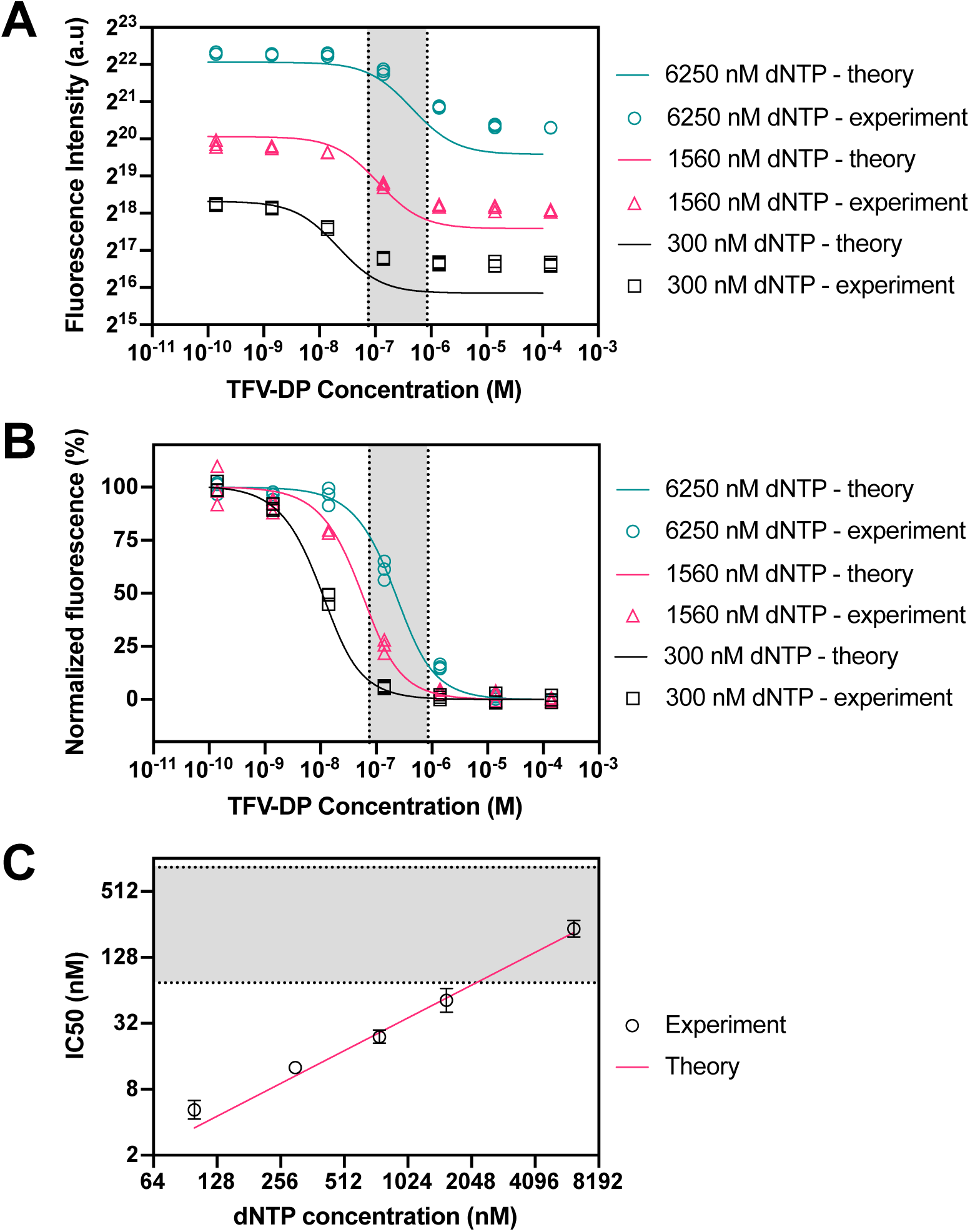
RESTRICT assay with TFV-DP in aqueous buffer at various dNTP concentrations. (A) RESTRICT assays with serial dilutions of TFV-DP at different dNTP concentrations (shown in the legend). Fluorescence intensity increases with dNTP concentration. Lines show fluorescence intensity calculated using the theory in Equations (1) to (7). (B) Normalized data showing that inhibition curves shift right towards higher TFV-DP as dNTP concentration increases. Lines show normalized fluorescence intensity values from the theory. (C) Graph of dNTP concentration versus IC_50_ values for experiments and theory. Grey shaded regions indicate the clinical range for TFV-DP adherence. N = 3, error bars indicate 95% confidence intervals.

We normalized the experimental data and theoretical calculations to more clearly visualize the effect of dNTP concentration on the position of the inhibition curves relative to the clinical range for TFV-DP adherence measurement. Figure 3B shows normalized endpoint RESTRICT assay fluorescence from experiments and theory. In both experiment and theory, as dNTP concentration increases the inhibition curves shift right, towards higher TFV-DP IC_50_ values. Figure 3C shows the measured IC_50_ values as a function of dNTP concentration along with a linear fit of IC_50_ values from theory. IC_50_ values increase linearly as a function of dNTP concentration. Figure 3B shows that the normalized experimental data is in better agreement with the model than the raw fluorescence data. This is because the model predicts the IC_50_ and response shape more accurately than it predicts end-point fluorescence, which depends on small variations in assay conditions and the plate reader response.

Using the theory described in equations (1) to (7) and the relationship between dNTP and IC_50_ in Figure 3C, we can design a RESTRICT assay with an IC_50_ value that enables us to differentiate TFV-DP concentrations that correspond to different adherence levels. To decrease the IC50, the dNTP and template concentration must decrease; however, there are limits to the relationship between IC_50_ and dNTP described in Figure 3C. For example, as the dNTP concentration is decreased to obtained lower IC_50_ values, template concentration must also be decreased to avoid dNTP depletion. Thus, at lower dNTP concentrations and corresponding lower template concentrations, the fluorescence intensity decreases (see Figure 3A). Thus, we can only decrease dNTP concentration until we run into the lower limit of detection of the fluorescence reader (see Figure 2C).

### RT activity assay in blood

TFV-DP accumulates intracellularly in both red blood cells and white blood cells. We chose to target TFV-DP in red blood cells because red blood cells are more abundant and are easier to lyse in order to release TFV-DP. As reported extensively in reverse transcription and DNA amplification assays, blood matrix components, such as immunoglobulins, inhibit RT activity and require sample preparation when working with whole blood.^56–58^ Autofluorescence from blood components^59^ can also obscure the signal from the RESTRICT assay.

We chose dilution in water as a simple and user-friendly strategy for sample preparation^60^ because it both lyses red blood cells and also reduces the concentration of blood matrix components thereby accomplishing our goals of simulating TFV-DP detection in a clinically relevant matrix and reducing unwanted suppression of reverse transcriptase activity.^60,61^ We tested various dilutions of whole blood in water to determine how much dilution was required to minimize non-specific inhibition. Figure 4A shows characterization of RT activity in diluted blood. The net fluorescence intensity, i.e. the difference between fluorescence from each RT activity assay with spiked blood and the background signal from no RT enzyme controls at each blood dilution, decreases as blood fraction increases and is indistinguishable from the background at 1.88% final concentration of blood. There is only a 20% decrease in fluorescence intensity compared to positive control (RT in buffer) at 0.25% final concentration of blood. Figure 4A shows that the non-specific inhibition of RT enzyme by blood matrix components also decreases with the concentration of blood in the assay.

**Figure 4.**
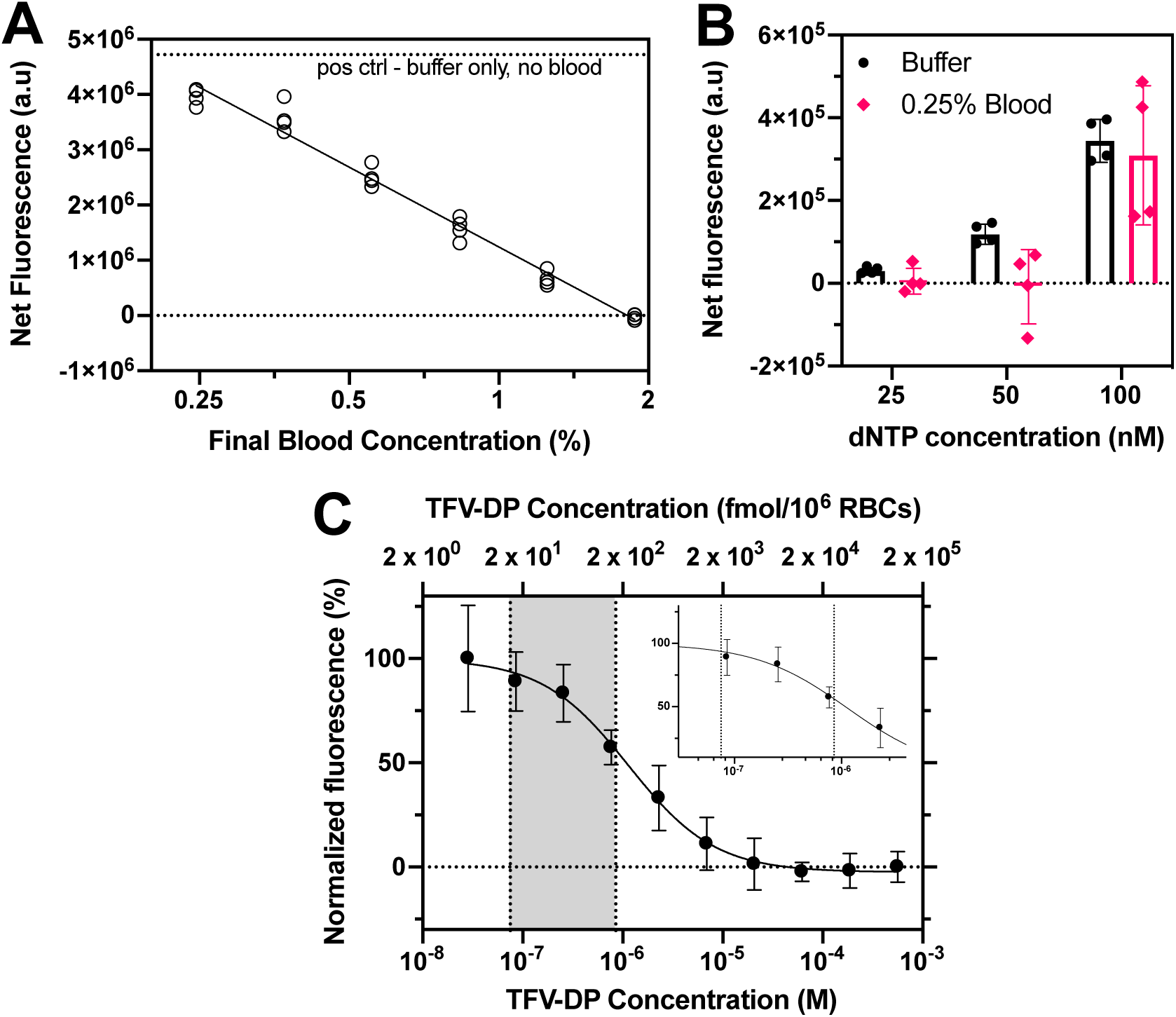
RT activity and inhibition assays in diluted blood. (A) RT activity assay with 500 nM dNTP and diluted whole blood spiked into the assay at various final concentrations to determine how much dilution was required to minimize non-specific RT inhibition by blood matrix components. (B) RT activity assay with 0.25% final blood concentration at low dNTP concentrations to determine the lowest dNTP concentration at which RT activity was detectable in blood. (C) Inhibition curve with TFV-DP spiked in diluted whole blood (0.25% blood final concentration) and 100 nM dNTP showing expected sigmoidal inhibition curve. Grey shaded region and inset show clinical range for TFV-DP adherence. N = 4, error bars indicate one standard deviation. The inset shows the same data of (C) over a narrower TFV-DP concentration range.

To perform RESTRICT assays in blood, we chose a blood dilution that reduces non-specific RT inhibition using Figure 4A. We also chose a dNTP concentration at which to run the assay to ensure that the clinical range of the assay overlapped with the linear region of the inhibition curve. There is a tradeoff between blood dilution and the required analytical sensitivity of the assay. Although greater blood dilution reduces non-specific inhibition of RT enzyme by blood matrix components (Figure 4A), greater dilution also decreases the available amount of analyte (TFV-DP) in blood.

Figures 3B and 3C show that the inhibition curve is shifted to lower TFV-DP concentrations by decreasing dNTP concentration. Figure 2C shows that the lowest template concentration and the lowest dNTP concentration at which we could detect RT activity in buffer was 0.25 nM template and 25 nM dNTP respectively. Anticipating that RT activity in blood would be more variable than in buffer, we re-ran RT activity assays in diluted blood to determine the lowest dNTP concentration at which we could perform RT activity assays. We chose a final concentration of 0.25% blood (dilution factor 400x) to minimize non-specific inhibition.

Figure 4B shows the net fluorescence measured in aqueous buffer and in 0.25% blood at dNTP concentrations of 25, 50, and 100 nM. Here the net fluorescence is the difference between the fluorescence measured from each data point minus the signal from a “no RT enzyme” control at the same conditions. This ensures that any variations in background signal from introducing diluted blood into the assay are accounted for. Figure 4B shows that there is a measurable fluorescence signal at 25 nM dNTP in buffer that increases gradually as the dNTP concentration is increased to 50 nM and 100 nM, consistent with Figure 2C. Conversely, the variation in RT activity when 0.25% blood is introduced means that the net fluorescence is zero at both 25 nM and 50 nM dNTP and a measurable signal from RT activity in 0.25% blood is only measurable at 100 nM. Thus, we determined that the lowest dNTP concentration that we could work with in 0.25% whole blood was 100 nM.

### RESTRICT assay in blood

Using the data in Figures 4A and 4B, we designed a RESTRICT assay for semi-quantitative measurement of clinically relevant TFV-DP concentrations spiked in diluted whole blood. Figure 4C shows data from an inhibition curve with 0.25% blood, 100 nM dNTP, at various TFV-DP concentrations around the clinical range for adherence measurement. We see the characteristic sigmoidal shape indicative of enzyme inhibition reactions.

We compared the normalized fluorescence intensities from the inhibition assay at low and high TFV-DP concentrations within the clinical range to determine whether we could distinguish concentrations that correspond to low and high adherence with statistical confidence. As shown in Table 1, the p-value is 0.013 for the unpaired t-test comparing fluorescence at 16.9 fmol/10^6^ RBCs TFV-DP, corresponding to low adherence, with the fluorescence at 152.3 fmol/10^6^ RBCs TFV-DP, indicating high adherence. These data demonstrate that the RESTRICT assay accurately distinguishes TFV-DP drug levels in spiked blood with high statistical confidence.

**Table 1.** Comparison between RESTRICT assay results at low and high concentrations within the clinical range for TFV-DP adherence measurement. N = 4.

|  |  |  |
| --- | --- | --- |
| TFV-DP Concentration (nM) | 85.4 | 768 |
| TFV-DP Concentration (fmol/10<sup>6</sup> RBCs) | 16.9 | 152.3 |
| Corresponding TFV-DP Adherence Level | Low | High |
| Normalized fluorescence (%) | 89.0 | 57.5 |
| 95% confidence interval | 66.4 – 111.5 | 44.1 – 70.6 |
| p-value for unpaired t-test | 0.013 |  |

## SUMMARY

In this paper, we present RESTRICT – an enzymatic assay for measurement of long-term PrEP and ART adherence. The assay measures cDNA formation by RT in the presence of TFV-DP. At high TFV-DP concentrations, cDNA chain termination occurs resulting in lower fluorescence signals from intercalating dye. We develop a probabilistic model for the RESTRICT assay by considering the likelihood of incorporation of TFV-DP into cDNA as a function of the relative concentrations of TFV-DP and dNTP. We show that decreasing dNTP concentration shifts RT inhibition curves to lower TFV-DP concentrations. The data and model show that there is a linear relationship between dNTP concentration and IC_50_ values. We spiked TFV-DP into hemolyzed whole blood and measured RT inhibition at TFV-DP concentrations within the clinical range for adherence measurement to demonstrate that the assay can operate in clinically relevant samples. The assay distinguished TFV-DP concentrations corresponding to low and high adherence suggesting that in future it might be a useful test to identify patients at risk of treatment failure.

Compared with ART and PrEP adherence monitoring by measuring TFV levels in urine, the RESTRICT assay measures TFV-DP and provides a longer-term measurement that is not susceptible to the white coat effect, where patients take their medications just before a doctor’s office visit. The RESTRICT assay can measure TFV-DP quickly, with less sample preparation, and capital investment than by LC/MS. The described assay gives measurable results within an hour whereas TFV-DP measurement by LC/MS requires several hours for sample processing and analysis and results are often not available for weeks because samples must be sent to one of the few centralized labs capable of measuring TFV-DP.

The RESTRICT assay has few liquid handling steps and relies on reagents and equipment (e.g. microplate reader and pipettes) that are commonly used and readily available in many clinical and biochemical laboratories. In addition, the cost per test is low since small reagent volumes (40 µL master mix) are required. The assay could be performed using fingerprick samples since < 5 µL of blood is required per assay and previous studies have established correlations between HIV drug metabolites in venous and fingerprick blood draws.^62^ The RESTRICT assay has the potential to be further integrated into a near-patient test for use in a doctor’s office or a patient’s home by replacing the microplate reader with a portable and inexpensive fluorescence reader or mobile phone^63,64^ and thus be a critical tool for preventing HIV transmission and drug resistance. We envision a test where assay reagents are lyophilized and stored in a tube or cartridge and the assay is initiated by adding blood and dilution buffer, incubating, and reading out results in a simple instrument.

The assay has some limitations that might preclude its use in particular contexts. For example, the assay does not yet distinguish between inhibition by NRTIs like TFV-DP and inhibition by non-nucleoside reverse transcriptase inhibitors (NNRTIs) like efavirenz that are sometimes included in ART regimens. For example, a patient may appear adherent if they took a recent dose of NNRTIs. This limitation is not a concern in PrEP clients since NNRTIs are not used in PrEP. Furthermore, as first line ART regimens in most of low- and middle-income countries switch from NNRTIs to dolutegravir (an integrase inhibitor),^65^ this limitation might not be a long-term concern.

The assay needs to be validated with clinical samples and benchmarked against drug level measurements by LC/MS. This work is innovative because it develops a new category of adherence measurement test that could allow patients and clinicians to monitor and improve long-term ART and PrEP adherence and healthcare outcomes.

## Supporting information

MATLAB Script

## SUPPORTING INFORMATION

Tables providing further additional details on the experimental conditions (master mix preparation for RESTRICT assays and TFV-DP dilutions in blood) are provided in the supplementary information.

## ACKNOWLEDGEMENTS

We are grateful for funding from the NIH (R01AI136648, R21AI127200, R01EB022630) and the M. J. Murdock Charitable Trust. Dr. Olanrewaju thanks the Mistletoe Research Foundation for funding and support. We are also grateful for helpful conversations with Jay Rutherford, Bob Atkinson, and Enos Kline at the University of Washington, as well as Dr Rebecca Sandlin and Dr Mehmet Toner at Harvard Medical School.

## REFERENCES

(1) UNAIDS Data 2019; UNAIDS, 2019; pp 1–476.

(2) WHO Expands Recommendation on Oral Pre-Exposure Prophylaxis of HIV Infection (PrEP); WHO/HIV/2015.48; World Health Organization (WHO), 2015.

(3) Castillo-Mancilla, J. R.; Haberer, J. E. Adherence Measurements in HIV: New Advancements in Pharmacologic Methods and Real-Time Monitoring. Curr. HIV/AIDS Rep. 2018, 15 (1), 49–59. https://doi.org/10.1007/s11904-018-0377-0.

(4) Abaasa, A.; Hendrix, C.; Gandhi, M.; Anderson, P.; Kamali, A.; Kibengo, F.; Sanders, E. J.; Mutua, G.; Bumpus, N. N.; Priddy, F.; et al. Utility of Different Adherence Measures for PrEP: Patterns and Incremental Value. AIDS Behav. 2018, 22 (4), 1165–1173. https://doi.org/10.1007/s10461-017-1951-y.

(5) Linnemayr, S.; Stecher, C. Behavioral Economics Matters for HIV Research: The Impact of Behavioral Biases on Adherence to Antiretrovirals (ARVs). AIDS Behav. 2015, 19 (11), 2069–2075. https://doi.org/10.1007/s10461-015-1076-0.

(6) Hamers, R. L.; Kityo, C.; Lange, J. M. A.; Wit, T. F. R. de; Mugyenyi, P. Global Threat from Drug Resistant HIV in Sub-Saharan Africa. BMJ 2012, 344, e4159. https://doi.org/10.1136/bmj.e4159.

(7) Mills, E. J.; Nachega, J. B.; Buchan, I.; Orbinski, J.; Attaran, A.; Singh, S.; Rachlis, B.; Wu, P.; Cooper, C.; Thabane, L.; et al. Adherence to Antiretroviral Therapy in Sub-Saharan Africa and North America: A Meta-Analysis. JAMA 2006, 296 (6), 679–690. https://doi.org/10.1001/jama.296.6.679.

(8) Ortego, C.; Huedo-Medina, T. B.; Llorca, J.; Sevilla, L.; Santos, P.; Rodríguez, E.; Warren, M. R.; Vejo, J. Adherence to Highly Active Antiretroviral Therapy (HAART): A Meta-Analysis. AIDS Behav. 2011, 15 (7), 1381–1396. https://doi.org/10.1007/s10461-011-9942-x.

(9) O’Connor, J. L.; Gardner, E. M.; Mannheimer, S. B.; Lifson, A. R.; Esser, S.; Telzak, E. E.; Phillips, A. N. Factors Associated With Adherence Amongst 5295 People Receiving Antiretroviral Therapy as Part of an International Trial. J. Infect. Dis. 2013, 208 (1), 40–49. https://doi.org/10.1093/infdis/jis731.

(10) Fonner, V. A.; Dalglish, S. L.; Kennedy, C. E.; Baggaley, R.; O’Reilly, K. R.; Koechlin, F. M.; Rodolph, M.; Hodges-Mameletzis, I.; Grant, R. M. Effectiveness and Safety of Oral HIV Preexposure Prophylaxis for All Populations. AIDS 2016, 30 (12), 1973. https://doi.org/10.1097/QAD.0000000000001145.

(11) Osterberg, L.; Blaschke, T. Adherence to Medication. N. Engl. J. Med. 2005, 353 (5), 487–497. https://doi.org/10.1056/NEJMra050100.

(12) Amico, K. R.; Stirratt, M. J. Adherence to Preexposure Prophylaxis: Current, Emerging, and Anticipated Bases of Evidence. Clin. Infect. Dis. 2014, 59 (Suppl_1), S55–S60. https://doi.org/10.1093/cid/ciu266.

(13) Simoni, J. M.; Kurth, A. E.; Pearson, C. R.; Pantalone, D. W.; Merrill, J. O.; Frick, P. A. Self-Report Measures of Antiretroviral Therapy Adherence: A Review with Recommendations for HIV Research and Clinical Management. AIDS Behav. 2006, 10 (3), 227–245. https://doi.org/10.1007/s10461-006-9078-6.

(14) McMahon, J. H.; Jordan, M. R.; Kelley, K.; Bertagnolio, S.; Hong, S. Y.; Wanke, C. A.; Lewin, S. R.; Elliott, J. H. Pharmacy Adherence Measures to Assess Adherence to Antiretroviral Therapy: Review of the Literature and Implications for Treatment Monitoring. Clin. Infect. Dis. 2011, 52 (4), 493–506. https://doi.org/10.1093/cid/ciq167.

(15) Haberer, J. E.; Kahane, J.; Kigozi, I.; Emenyonu, N.; Hunt, P.; Martin, J.; Bangsberg, D. R. Real-Time Adherence Monitoring for HIV Antiretroviral Therapy. AIDS Behav. 2010, 14 (6), 1340–1346. https://doi.org/10.1007/s10461-010-9799-4.

(16) Garrison, L. E.; Haberer, J. E. Technological Methods to Measure Adherence to Antiretroviral Therapy and Preexposure Prophylaxis: Curr. Opin. HIV AIDS 2017, 12 (5), 467–474. https://doi.org/10.1097/COH.0000000000000393.

(17) Chai, P. R.; Castillo-Mancilla, J.; Buffkin, E.; Darling, C.; Rosen, R. K.; Horvath, K. J.; Boudreaux, E. D.; Robbins, G. K.; Hibberd, P. L.; Boyer, E. W. Utilizing an Ingestible Biosensor to Assess Real-Time Medication Adherence. J. Med. Toxicol. 2015, 11 (4), 439–444. https://doi.org/10.1007/s13181-015-0494-8.

(18) Frasca, K.; Morrow, M.; Coyle, R. P.; Coleman, S. S.; Ellison, L.; Bushman, L. R.; Kiser, J. J.; Zheng, J.-H.; Mawhinney, S.; Anderson, P. L.; et al. Emtricitabine Triphosphate in Dried Blood Spots Is a Predictor of Viral Suppression in HIV Infection and Reflects Short-Term Adherence to Antiretroviral Therapy. J. Antimicrob. Chemother. 2019, 74 (5), 1395–1401. https://doi.org/10.1093/jac/dky559.

(19) Castillo-Mancilla, J.; Coyle, R.; Coleman, S.; Morrow, M.; Gardner, E.; Zheng, J.-H.; Ellison, L.; Bushman, L. R.; Kiser, J.; MaWhinney, S.; et al. Cascade of ART Adherence in Virologically-Suppressed Persons Living with HIV. AIDS Res. Hum. Retroviruses 2019. https://doi.org/10.1089/AID.2019.0024.

(20) Coyle, R. P.; Schneck, C. D.; Morrow, M.; Coleman, S. S.; Gardner, E. M.; Zheng, J.-H.; Ellison, L.; Bushman, L. R.; Kiser, J. J.; Mawhinney, S.; et al. Engagement in Mental Health Care Is Associated with Higher Cumulative Drug Exposure and Adherence to Antiretroviral Therapy. AIDS Behav. 2019. https://doi.org/10.1007/s10461-019-02441-8.

(21) Morrow, M.; MaWhinney, S.; Coyle, R. P.; Coleman, S. S.; Gardner, E. M.; Zheng, J.-H.; Ellison, L.; Bushman, L. R.; Kiser, J. J.; Anderson, P. L.; et al. Predictive Value of Tenofovir Diphosphate in Dried Blood Spots for Future Viremia in Persons Living With HIV. J. Infect. Dis. 2019, 220 (4), 635–642. https://doi.org/10.1093/infdis/jiz144.

(22) Organization, W. H.; UNAIDS. Antiretroviral Medicines in Low- and Middle-Income Countries: Forecasts of Global and Regional Demand for 2013-2016; World Health Organization, 2014.

(23) Kearney, B. P.; Flaherty, J. F.; Shah, J. Tenofovir Disoproxil Fumarate. Clin. Pharmacokinet. 2004, 43 (9), 595–612. https://doi.org/10.2165/00003088-200443090-00003.

(24) Castillo-Mancilla, J. R.; Zheng, J.-H.; Rower, J. E.; Meditz, A.; Gardner, E. M.; Predhomme, J.; Fernandez, C.; Langness, J.; Kiser, J. J.; Bushman, L. R.; et al. Tenofovir, Emtricitabine, and Tenofovir Diphosphate in Dried Blood Spots for Determining Recent and Cumulative Drug Exposure. AIDS Res. Hum. Retroviruses 2012, 121010062750004. https://doi.org/10.1089/aid.2012.0089.

(25) Anderson, P. L. What Can Urine Tell Us About Medication Adherence? EClinicalMedicine 2018, 2, 5–6. https://doi.org/10.1016/j.eclinm.2018.09.003.

(26) Spinelli, M. A.; Glidden, D. V.; Anderson, P. L.; Gandhi, M.; Cohen, S.; Vittinghoff, E.; Coleman, M. E.; Scott, H.; Bacon, O.; Elion, R.; et al. Short-Term Adherence Marker to PrEP Predicts Future Nonretention in a Large PrEP Demo Project: Implications for Point-of-Care Adherence Testing. J Acquir Immune Defic Syndr 2019, 81 (2), 5.

(27) Anderson, P. L.; Liu, A. Y.; Castillo-Mancilla, J. R.; Gardner, E. M.; Seifert, S. M.; McHugh, C.; Wagner, T.; Campbell, K.; Morrow, M.; Ibrahim, M.; et al. Intracellular Tenofovir-Diphosphate and Emtricitabine-Triphosphate in Dried Blood Spots Following Directly Observed Therapy. Antimicrob. Agents Chemother. 2018, 62 (1), e01710–17. https://doi.org/10.1128/AAC.01710-17.

(28) Castillo-Mancilla, J. R.; Morrow, M.; Coyle, R. P.; Coleman, S. S.; Gardner, E. M.; Zheng, J.-H.; Ellison, L.; Bushman, L. R.; Kiser, J. J.; Mawhinney, S.; et al. Tenofovir Diphosphate in Dried Blood Spots Is Strongly Associated With Viral Suppression in Individuals With Human Immunodeficiency Virus Infections. Clin. Infect. Dis. 2019, 68 (8), 1335–1342. https://doi.org/10.1093/cid/ciy708.

(29) Grant, R. M.; Anderson, P. L.; McMahan, V.; Liu, A.; Amico, K. R.; Mehrotra, M.; Hosek, S.; Mosquera, C.; Casapia, M.; Montoya, O.; et al. Uptake of Pre-Exposure Prophylaxis, Sexual Practices, and HIV Incidence in Men and Transgender Women Who Have Sex with Men: A Cohort Study. Lancet Infect. Dis. 2014, 14 (9), 820–829. https://doi.org/10.1016/S1473-3099(14)70847-3.

(30) Pratt, G. W.; Fan, A.; Melakeberhan, B.; Klapperich, C. M. A Competitive Lateral Flow Assay for the Detection of Tenofovir. Anal. Chim. Acta 2018, 1017, 34–40. https://doi.org/10.1016/j.aca.2018.02.039.

(31) Gandhi, M.; Bacchetti, P.; Rodrigues, W. C.; Spinelli, M.; Koss, C. A.; Drain, P. K.; Baeten, J. M.; Mugo, N. R.; Ngure, K.; Benet, L. Z.; et al. Development and Validation of an Immunoassay for Tenofovir in Urine as a Real-Time Metric of Antiretroviral Adherence. EClinicalMedicine 2018, 2–3, 22–28. https://doi.org/10.1016/j.eclinm.2018.08.004.

(32) Spinelli, M. A.; Glidden, D. V.; Rodrigues, W. C.; Wang, G.; Vincent, M.; Okochi, H.; Kuncze, K.; Mehrotra, M.; Defechereux, P.; Buchbinder, S. P.; et al. Low Tenofovir Level in Urine by a Novel Immunoassay Is Associated with Seroconversion in a Preexposure Prophylaxis Demonstration Project: AIDS 2019, 33 (5), 867–872. https://doi.org/10.1097/QAD.0000000000002135.

(33) Koenig, H. C.; Mounzer, K.; Daughtridge, G. W.; Sloan, C. E.; Lalley-Chareczko, L.; Moorthy, G. S.; Conyngham, S. C.; Zuppa, A. F.; Montaner, L. J.; Tebas, P. Urine Assay for Tenofovir to Monitor Adherence in Real Time to Tenofovir Disoproxil Fumarate/Emtricitabine as Pre-Exposure Prophylaxis. HIV Med. 2017, 18 (6), 412–418. https://doi.org/10.1111/hiv.12518.

(34) Gandhi, M. M.; Bacchetti, P.; Spinelli, M. A. M. a; Okochi, H.; Baeten, J. M.; Siriprakaisil, O. M. e; Klinbuayaem, V. M. e; Rodrigues, W. C. M. f; Wang, G.; Vincent, M. M. f; et al. Validation of a Urine Tenofovir Immunoassay for Adherence Monitoring to PrEP and ART and Establishing the Cutoff for a Point-of-Care Test. J. Acquir. Immune Defic. Syndr. 2019, 81 (1), 72–77. https://doi.org/10.1097/QAI.0000000000001971.

(35) Anderson, P. L.; Glidden, D. V.; Liu, A.; Buchbinder, S.; Lama, J. R.; Guanira, J. V.; McMahan, V.; Bushman, L. R.; Casapía, M.; Montoya-Herrera, O.; et al. Emtricitabine-Tenofovir Concentrations and Pre-Exposure Prophylaxis Efficacy in Men Who Have Sex with Men. Sci. Transl. Med. 2012, 4 (151), 151ra125–151ra125. https://doi.org/10.1126/scitranslmed.3004006.

(36) Porstmann, T.; Meissner, K.; Glaser, R.; Döpel, S.-H.; Sydow, G. A Sensitive Non-Isotopic Assay Specific for HIV-1 Associated Reverse Transcriptase. J. Virol. Methods 1991, 31 (2), 181–188. https://doi.org/10.1016/0166-0934(91)90156-T.

(37) Suzuki, K.; Craddock, B. P.; Okamoto, N.; Kano, T.; Steigbigel, R. T. Poly A-Linked Colorimetric Microtiter Plate Assay for HIV Reverse Transcriptase. J. Virol. Methods 1993, 44 (2), 189–198. https://doi.org/10.1016/0166-0934(93)90054-U.

(38) Suzuki, K.; Craddock, B. P.; Kano, T.; Steigbigel, R. T. Chemiluminescent Enzyme-Linked Immunoassay for Reverse Transcriptase, Illustrated by Detection of HIV Reverse Transcriptase. Anal. Biochem. 1993, 210 (2), 277–281. https://doi.org/10.1006/abio.1993.1196.

(39) Balzarini, J.; De Clercq, E. [25] Analysis of Inhibition of Retroviral Reverse Transcriptase. In Methods in Enzymology; Viral Polymerases and Related Proteins; Academic Press, 1996; Vol. 275, pp 472–502. https://doi.org/10.1016/S0076-6879(96)75027-9.

(40) Arnold, B. A.; Hepler, R. W.; Keller, P. M. One-Step Fluorescent Probe Product-Enhanced Reverse Transcriptase Assay. BioTechniques 1998, 25 (1), 98–106.

(41) Balzarini, J.; Pérez-Pérez, M. J.; San-Félix, A.; Camarasa, M. J.; Bathurst, I. C.; Barr, P. J.; Clercq, E. D. Kinetics of Inhibition of Human Immunodeficiency Virus Type 1 (HIV-1) Reverse Transcriptase by the Novel HIV-1-Specific Nucleoside Analogue [2’,5’-Bis-O-(Tert-Butyldimethylsilyl)-Beta-D-Ribofuranosyl]-3’-Spiro-5 “-(4”-Amino-1”,2”-Oxathiole-2”,2”-Dioxide)Thymine (TSAO-T). J. Biol. Chem. 1992, 267 (17), 11831–11838.

(42) Cheng, Y. C.; Dutschman, G. E.; Bastow, K. F.; Sarngadharan, M. G.; Ting, R. Y. Human Immunodeficiency Virus Reverse Transcriptase. General Properties and Its Interactions with Nucleoside Triphosphate Analogs. J. Biol. Chem. 1987, 262 (5), 2187–2189.

(43) Kokkula, C.; Palanisamy, N.; Ericstam, M.; Lennerstrand, J. SYBR Green II Dye-Based Real-Time Assay for Measuring Inhibitor Activity Against HIV-1 Reverse Transcriptase. Mol. Biotechnol. 2016, 58 (10), 619–625. https://doi.org/10.1007/s12033-016-9961-y.

(44) Frezza, C.; Balestrieri, E.; Marino-Merlo, F.; Mastino, A.; Macchi, B. A Novel, Cell-Free PCR-Based Assay for Evaluating the Inhibitory Activity of Antiretroviral Compounds against HIV Reverse Transcriptase. J. Med. Virol. 2014, 86 (1), 1–7. https://doi.org/10.1002/jmv.23748.

(45) Marino-Merlo, F.; Frezza, C.; Papaianni, E.; Valletta, E.; Mastino, A.; Macchi, B. Development and Evaluation of a Simple and Effective RT-QPCR Inhibitory Assay for Detection of the Efficacy of Compounds towards HIV Reverse Transcriptase. Appl. Microbiol. Biotechnol. 2017, 101 (22), 8249–8258. https://doi.org/10.1007/s00253-017-8544-6.

(46) Boyer, P. L.; Tantillo, C.; Jacobo-Molina, A.; Nanni, R. G.; Ding, J.; Arnold, E.; Hughes, S. H. Sensitivity of Wild-Type Human Immunodeficiency Virus Type 1 Reverse Transcriptase to Dideoxynucleotides Depends on Template Length; the Sensitivity of Drug-Resistant Mutants Does Not. Proc. Natl. Acad. Sci. 1994, 91 (11), 4882–4886. https://doi.org/10.1073/pnas.91.11.4882.

(47) Boyer, P. L.; Clark, P. K.; Hughes, S. H. HIV-1 and HIV-2 Reverse Transcriptases: Different Mechanisms of Resistance to Nucleoside Reverse Transcriptase Inhibitors. J. Virol. 2012, 86 (10), 5885–5894. https://doi.org/10.1128/JVI.06597-11.

(48) Marino-Merlo, F.; Macchi, B.; Armenia, D.; Bellocchi, M. C.; Ceccherini-Silberstein, F.; Mastino, A.; Grelli, S. Focus on Recently Developed Assays for Detection of Resistance/Sensitivity to Reverse Transcriptase Inhibitors. Appl. Microbiol. Biotechnol. 2018. https://doi.org/10.1007/s00253-018-9390-x.

(49) Bebenek, K.; Abbotts, J.; Roberts, J. D.; Wilson, S. H.; Kunkel, T. A. Specificity and Mechanism of Error-Prone Replication by Human Immunodeficiency Virus-1 Reverse Transcriptase. J. Biol. Chem. 1989, 264 (28), 16948–16956.

(50) Dragan, A. I.; Casas-Finet, J. R.; Bishop, E. S.; Strouse, R. J.; Schenerman, M. A.; Geddes, C. D. Characterization of PicoGreen Interaction with DsDNA and the Origin of Its Fluorescence Enhancement upon Binding. Biophys. J. 2010, 99 (9), 3010–3019. https://doi.org/10.1016/j.bpj.2010.09.012.

(51) Singer, V. L.; Jones, L. J.; Yue, S. T.; Haugland, R. P. Characterization of PicoGreen Reagent and Development of a Fluorescence-Based Solution Assay for Double-Stranded DNA Quantitation. Anal. Biochem. 1997, 249 (2), 228–238. https://doi.org/10.1006/abio.1997.2177.

(52) Hoffman, A. D.; Banapour, B.; Levy, J. A. Characterization of the AIDS-Associated Retrovirus Reverse Transcriptase and Optimal Conditions for Its Detection in Virions. Virology 1985, 147 (2), 326–335. https://doi.org/10.1016/0042-6822(85)90135-7.

(53) Zadeh, J. N.; Steenberg, C. D.; Bois, J. S.; Wolfe, B. R.; Pierce, M. B.; Khan, A. R.; Dirks, R. M.; Pierce, N. A. NUPACK: Analysis and Design of Nucleic Acid Systems. J. Comput. Chem. 2011, 32 (1), 170–173. https://doi.org/10.1002/jcc.21596.

(54) Huber, H. E.; McCoy, J. M.; Seehra, J. S.; Richardson, C. C. Human Immunodeficiency Virus 1 Reverse Transcriptase. Template Binding, Processivity, Strand Displacement Synthesis, and Template Switching. J. Biol. Chem. 1989, 264 (8), 4669–4678.

(55) Brooks, H. B.; Geeganage, S.; Kahl, S. D.; Montrose, C.; Sittampalam, S.; Smith, M. C.; Weidner, J. R. Basics of Enzymatic Assays for HTS. In Assay Guidance Manual; Sittampalam, G. S., Coussens, N. P., Brimacombe, K., Grossman, A., Arkin, M., Auld, D., Austin, C., Baell, J., Bejcek, B., Chung, T. D. Y., et al., Eds.; Eli Lilly & Company and the National Center for Advancing Translational Sciences: Bethesda (MD), 2004.

(56) Sidstedt, M.; Hedman, J.; Romsos, E. L.; Waitara, L.; Wadsö, L.; Steffen, C. R.; Vallone, P. M.; Rådström, P. Inhibition Mechanisms of Hemoglobin, Immunoglobulin G, and Whole Blood in Digital and Real-Time PCR. Anal. Bioanal. Chem. 2018, 410 (10), 2569–2583. https://doi.org/10.1007/s00216-018-0931-z.

(57) Al-Soud, W. A.; Rådström, P. Purification and Characterization of PCR-Inhibitory Components in Blood Cells. J. Clin. Microbiol. 2001, 39 (2), 485–493. https://doi.org/10.1128/JCM.39.2.485-493.2001.

(58) Al-Soud, W. A.; Jönsson, L. J.; Rådström, P. Identification and Characterization of Immunoglobulin G in Blood as a Major Inhibitor of Diagnostic PCR. J. Clin. Microbiol. 2000, 38 (1), 345–350.

(59) Hoffman, R. A.; Hansen, W. P. Immunofluorescent Analysis of Blood Cells by Flow Cytometry. Int. J. Immunopharmacol. 1981, 3 (3), 249–254. https://doi.org/10.1016/0192-0561(81)90018-7.

(60) Cai, D.; Behrmann, O.; Hufert, F.; Dame, G.; Urban, G. Direct DNA and RNA Detection from Large Volumes of Whole Human Blood. Sci. Rep. 2018, 8 (1), 3410. https://doi.org/10.1038/s41598-018-21224-0.

(61) Higuchi, R. Simple and Rapid Preparation of Samples for PCR. In PCR Technology: Principles and Applications for DNA Amplification; Erlich, H. A., Ed.; Palgrave Macmillan UK: London, 1989; pp 31–38. https://doi.org/10.1007/978-1-349-20235-5_4.

(62) Castillo-Mancilla, J.; Seifert, S.; Campbell, K.; Coleman, S.; McAllister, K.; Zheng, J.-H.; Gardner, E. M.; Liu, A.; Glidden, D. V.; Grant, R.; et al. Emtricitabine-Triphosphate in Dried Blood Spots as a Marker of Recent Dosing. Antimicrob. Agents Chemother. 2016, 60 (11), 6692–6697. https://doi.org/10.1128/AAC.01017-16.

(63) Kim, D.; Wei, Q.; Kim, D. H.; Tseng, D.; Zhang, J.; Pan, E.; Garner, O.; Ozcan, A.; Di Carlo, D. Enzyme-Free Nucleic Acid Amplification Assay Using a Cellphone-Based Well Plate Fluorescence Reader. Anal. Chem. 2018, 90 (1), 690–695. https://doi.org/10.1021/acs.analchem.7b03848.

(64) Mak, W. C.; Beni, V.; Turner, A. P. F. Lateral-Flow Technology: From Visual to Instrumental. TrAC Trends Anal. Chem. 2016, 79, 297–305. https://doi.org/10.1016/j.trac.2015.10.017.

(65) Dorward, J.; Lessells, R.; Drain, P. K.; Naidoo, K.; de Oliveira, T.; Pillay, Y.; Abdool Karim, S. S.; Garrett, N. Dolutegravir for First-Line Antiretroviral Therapy in Low-Income and Middle-Income Countries: Uncertainties and Opportunities for Implementation and Research. Lancet HIV 2018, 5 (7), e400–e404. https://doi.org/10.1016/S2352-3018(18)30093-6.

