## Supplementary Information for "An enzymatic assay to measure long-term adherence to pre-exposure prophylaxis and antiretroviral therapy"

Jonathan D. Posner^1,8,9^

**Corresponding Authors:**

Jonathan D. Posner:

Paul K. Drain:

**Affiliations:**

1. Department of Mechanical Engineering, University of Washington, Seattle, USA
2. Center for Engineering in Medicine, Massachusetts General Hospital, Harvard Medical School, Boston
3. Department of Materials Science & Engineering, University of Washington, Seattle
4. Independent Consultant
5. Department of Epidemiology, University of Washington, Seattle
6. Department of Global Health, University of Washington, Seattle
7. Department of Medicine, University of Washington, Seattle
8. Department of Chemical Engineering, University of Washington, Seattle
9. Department of Family Medicine, University of Washington, Seattle

**Abstract:**

The supporting information shows:

1. Master mixes for RESTRICT assays in buffer.
2. Tenofovir diphosphate dilutions in blood.
1. **Master mixes for RESTRICT assays in buffer**

As described in the Experimental Section of the manuscript, master mixes were prepared to optimize RT activity and ensure that the REverse TranscrIptase Chain Termination (RESTRICT) assay operates in a regime that provides high assay signal in the absence of nucleotide reverse transcriptase inhibitors (NRTIs). Master mixes for the RESTRICT assay with TFV-DP consisted of DNA template, primer, dNTP solution, TFV‑DP, and HIV-1 RT as summarized in Table 1 below. Serial dilutions of TFV-DP in buffer spanning a concentration range of 1 – 10,000 nM were prepared to generate curves representative of TFV-DP inhibition.

Table S1: Volumes and concentrations of reagents used to prepare master mixes for RESTRICT assay.

| **Reagent** | **Stock Concentration [M]** | **Volume per reaction (µL)** | **# Reactions** | **Total Volume (µL)** | **Stock volume (µL)** | **Buffer volume (µL)** | **Final Concentration [M]** |
| --- | --- | --- | --- | --- | --- | --- | --- |
| TCAA template | 8.00E-07 | 5 | 29.7 | 148.5 | 3.7 | 144.8 | 2.5E-09 |
| 16S rRNA primer | 8.00E-07 | 5 | 29.7 | 148.5 | 37.1 | 111.4 | 2.5E-08 |
| dNTP | 5.00E-04 | 20 | 29.7 | 594 | 3.7 | 590.3 | 1.6E-06 |
|  |  | **30** | **29.7** | **891** |  |  |  |
| TFV-DP in buffer | varies | 5 | 29.7 | 148.5 |  |  |  |
| HIV-1 RT | 8.55E-06 | 5 | 26.4 | 132 | 12.4 | 119.6 | 1.0E-07 |
|  |  | **40** |  | **1171.5** |  |  |  |
| **After assay** |  |  |  |  |  |  |  |
| Picogreen | 400 | 40 | 29.7 | 1188 | 5.9 | 1182.1 | 1 |

2. **Tenofovir diphosphate dilutions in blood**

We performed RESTRICT assays with TFV‑DP in whole blood to demonstrate that the RESTRICT assay could operate in a clinically relevant sample and at concentrations relevant for TFV-DP adherence monitoring. We used dilution in water as a simple sample preparation strategy to lyse red blood cells and reduce non-specific inhibition by blood matrix components.^1,2^ To simulate clinical samples, we spiked TFV‑DP into the diluted whole blood at concentrations that correspond to the clinical range for TFV‑DP adherence. The median concentration of TFV‑DP in red blood cells ranges from 15 – 170 fmol/10^6^ RBCs.^3^ Assuming an average hematocrit of 40% and an average RBC count of $5\times{10}^{6}$ RBCs/µL for adults, the clinical range for TFV‑DP adherence corresponds to 75 – 850nM TFV‑DP in whole blood.

To prepare the diluted whole blood at 2% final concentration, we added 13.3 µL of whole blood was added to 661.7 µL of nuclease‑free water. Next, we added 4.6 µL of 110 µM TFV‑DP in RT assay buffer to 40.4 µL of the 2% blood to obtain a solution with 11 µM of TFV‑DP in 2% blood. We further diluted the TFV-DP in blood by adding 15 µL of 11 µM TFV‑DP in 2% blood to 30 µL of 2% blood. Eight additional 1 in 3 dilutions of TFV‑DP in 2% blood were carried as summarized in Table S2. TFV-DP concentrations in the assay were chosen so that the corresponding concentration of TFV‑DP in undiluted blood (i.e. multiplied by 400 to account for 50X dilution of blood and 8X dilution of TFV‑DP in master mix) spanned a range of 5.7 – 110, 000 fmol/10^6^ RBCs and thus covers the clinical range for TFV‑DP adherence.

Table S2: Serial dilutions of TFV‑DP in 2% whole blood.

| **Tube #** | **Stock Concentration (M)** | **Volume per reaction (µL)** | **# Reactions** | **Total Volume (µL)** | **Stock volume (µL)** | **Diluted Blood Volume (µL)** | **Final Concentration in tube [M]** | **Final concentration in assay [M]** | **Corresponding concentration in undiluted whole blood**  **(i.e. 400X)**  **[M]** | **Corresponding concentration in undiluted whole blood**  **(i.e. 400X) [fmol/10^6^ RBCs]** |
| --- | --- | --- | --- | --- | --- | --- | --- | --- | --- | --- |
| 1 | 1.1E-04 | 5 | 9 | 45 | 4.6 | 40.4 | 1.1E-05 | 1.4E-06 | 5.6E-04 | 1.1E+05 |
| 2 | 1.1E-05 | 5 | 9 | 45 | 15.0 | 30.0 | 3.7E-06 | 4.7E-07 | 1.9E-04 | 3.7E+04 |
| 3 | 3.7E-06 | 5 | 9 | 45 | 15.0 | 30.0 | 1.2E-06 | 1.6E-07 | 6.2E-05 | 1.2E+04 |
| 4 | 1.2E-06 | 5 | 9 | 45 | 15.0 | 30.0 | 4.1E-07 | 5.2E-08 | 2.1E-05 | 4.1E+03 |
| 5 | 4.1E-07 | 5 | 9 | 45 | 15.0 | 30.0 | 1.4E-07 | 1.7E-08 | 6.9E-06 | 1.4E+03 |
| 6 | 1.4E-07 | 5 | 9 | 45 | 15.0 | 30.0 | 4.6E-08 | 5.8E-09 | 2.3E-06 | 4.6E+02 |
| 7 | 4.6E-08 | 5 | 9 | 45 | 15.0 | 30.0 | 1.5E-08 | 1.9E-09 | 7.7E-07 | 1.5E+02 |
| 8 | 1.5E-08 | 5 | 9 | 45 | 15.0 | 30.0 | 5.1E-09 | 6.4E-10 | 2.6E-07 | 5.1E+01 |
| 9 | 5.1E-09 | 5 | 9 | 45 | 15.0 | 30.0 | 1.7E-09 | 2.1E-10 | 8.5E-08 | 1.7E+01 |
| 10 | 1.7E-09 | 5 | 9 | 45 | 15.0 | 30.0 | 5.7E-10 | 7.1E-11 | 2.8E-08 | 5.7E+00 |

**REFERENCES**

(1)	Higuchi, R. Simple and Rapid Preparation of Samples for PCR. In *PCR Technology: Principles and Applications for DNA Amplification*; Erlich, H. A., Ed.; Palgrave Macmillan UK: London, 1989; pp 31–38. https://doi.org/10.1007/978-1-349-20235-5_4.

(2)	Cai, D.; Behrmann, O.; Hufert, F.; Dame, G.; Urban, G. Direct DNA and RNA Detection from Large Volumes of Whole Human Blood. *Sci. Rep.* **2018**, *8* (1), 3410. https://doi.org/10.1038/s41598-018-21224-0.

(3)	Castillo-Mancilla, J. R.; Zheng, J.-H.; Rower, J. E.; Meditz, A.; Gardner, E. M.; Predhomme, J.; Fernandez, C.; Langness, J.; Kiser, J. J.; Bushman, L. R.; et al. Tenofovir, Emtricitabine, and Tenofovir Diphosphate in Dried Blood Spots for Determining Recent and Cumulative Drug Exposure. *AIDS Res. Hum. Retroviruses* **2012**, 121010062750004. https://doi.org/10.1089/aid.2012.0089.
