## Supplementary material for "An enzymatic assay to measure long-term adherence to pre-exposure prophylaxis and antiretroviral therapy": MATLAB Script: Olanrewaju_etal_MATLAB_Script.m

```
close all
clc
%clear all

%%%%% INPUT DNA STRANDS
TTCA = "TTCATTCATTCATTCATTCATTCATTCATTCATTCATTCATTCATTCATTCATTCATTCATTCATTCATTCATTCATTCATTCATTCATTCATTCATTCATTCATTCATTCATTCATTCATTCATTCATTCATTCATTCATTCATTCATTCATTCATTCATTCATTCATTCATTCATTCA";
DNAstrand = [TTCA];

%%%%% INPUT REAGENT CONCENTRATIONS
dNTP_Conc = 100*10^-9; 
Inhibitor_Conc = logspace(-10,-4,1000);
Inhibitor_Conc_nM = 1e9 * Inhibitor_Conc;
Logspace = Inhibitor_Conc(2)-Inhibitor_Conc(1);

%%%%% INPUT RELATIVE DYE AFFINITIES
dyedsDNA = 1;
dyessDNA = 1/11;
templateConcentration = dNTP_Conc/400; %ratio of dNTP to template concentration is 400 to 1

%%%%% INPUT RELATIVE INHIBITOR AFFINITY
K_aff = 0.5;
Tcount_matrix = count(DNAstrand, "T");
Length_matrix = strlength(DNAstrand);

%%%%% CALCULATIONS START HERE
for k = 1:length(DNAstrand)
    DNAstrand = DNAstrand(k);
    DNAlength = strlength(DNAstrand);
    Tlocation = strfind(DNAstrand,"T");
    Tcount = count(DNAstrand,"T");
    %%%%% CALCULATESPRODUCTS
    for j = 1:length(Inhibitor_Conc)
        Initial = 1; %% Set initial value to 1, since you have 100% ssDNA templates at the beginning
        for i = 1:Tcount
            Completion(i) = (dNTP_Conc/(K_aff*Inhibitor_Conc(j)+dNTP_Conc))^i;
            dsDNA_Fragment_Probability(i) = Initial - Completion(i);
            Initial = Completion(i);
        end
        %%%%% CALCULATES COMPLETED PRODUCTS (dsDNA)
        
        dsDNAFull(j) = dyedsDNA * Completion(end) * DNAlength * templateConcentration;
        dsDNAFragment(j) = dyedsDNA * sum(dsDNA_Fragment_Probability.*Tlocation) * templateConcentration;
        totaldsDNA(j) = dsDNAFull(j) + dsDNAFragment(j);
        
        %%%%% CALCULATES UNFINISHED PRODUCTS (ssDNA)
        totalssDNA(j) = dyessDNA * sum(dsDNA_Fragment_Probability.*(DNAlength-Tlocation)) * templateConcentration;
    end
    
% Calculate total signal including both dsDNA and ssDNA

    %%%%% CALCULATES TOTAL SIGNAL (INCLUDES BOTH dsDNA and ssDNA)
    %% Primer is 20 nt long
    primerLength = 20;
    %% Primer concentration = 10X template concentration 
    %% Assuming steady state reaction that uses up all the DNA template, then fluorescence due to dsDNA from bound primer can be calculated as:
    boundPrimerFluorescence = primerLength * dyedsDNA * ones(size(Inhibitor_Conc)) * templateConcentration; 
    %% Calculate primerFluorescence = fluorescence from unused primer by multiplying primer concentration, primer length, and fluorescence from ssDNA
    excessPrimerConcentration = 9 * templateConcentration;
    primerFluorescence = excessPrimerConcentration*primerLength*dyessDNA*ones(size(Inhibitor_Conc)); 
    dsDNAFullWithPrimer = dsDNAFull + primerFluorescence;
    totaldsDNAWithPrimer = totaldsDNA + primerFluorescence;
    TotalProduct = totalssDNA + totaldsDNAWithPrimer + boundPrimerFluorescence;
    TotalProduct_dsDNAOnly = dyedsDNA*totaldsDNA + boundPrimerFluorescence;
    TotalProduct_WithPrimer = dyessDNA*totalssDNA + dyedsDNA*totaldsDNA + primerFluorescence + boundPrimerFluorescence;
    
    clearvars Initial Completion Total dsDNA_Fragment_Probability
end

figure
semilogx(Inhibitor_Conc,(dsDNAFull),'k-.')
hold on
%semilogx(Inhibitor_Conc,dsDNAFragment,'k--')
%hold on
semilogx(Inhibitor_Conc,(totaldsDNA),'g-')
hold on
semilogx(Inhibitor_Conc,(totalssDNA),'b-')
hold on
semilogx(Inhibitor_Conc,(primerFluorescence),'b--')
hold on
semilogx(Inhibitor_Conc,(TotalProduct),'r-')
hold on
legend('Fluorescence from full-length cDNA only','Fluorescence from full-length and fragment cDNA','Unused ssDNA template','Excess Primer','Total Fluorescence')
xlabel('TFV-DP Concentration [M]')
ylabel('Fluorescence intensity (a.u)')
% Normalize theoretical data
% $${\textrm{Theoretical}}_{\textrm{normalized}} =100*\;\frac{{\textrm{Theoretical}}_i 
% -{\textrm{Theoretical}}_{\min } }{{\textrm{Theoretical}}_{\max } -{\textrm{Expeirmental}}_{\min 
% } }\;%$$

%%%%% NORMALIZATION PROCEDURE -- Total Product
for i = 1:length(TotalProduct)
    TotalNormalized(i) = 100*(TotalProduct(i)-min(TotalProduct))/(max(TotalProduct)-min(TotalProduct));
end

%% Normalized data - full length and fragment cDNA only, no ssDNA
for i = 1:length(TotalProduct)
    TotalNormalized_dsDNAOnly(i) = 100*(totaldsDNA(i)-min(totaldsDNA))/(max(totaldsDNA)-min(totaldsDNA));
end

%% Normalized data - full length cDNA only
for i = 1:length(totaldsDNA)
    TotalNormalized_dsDNAFullProduct(i) = 100*(dsDNAFull(i)-min(dsDNAFull))/(max(dsDNAFull)-min(dsDNAFull));
end

%% Normalized data - with Primer contributions
for i = 1:length(TotalProduct)
    TotalNormalized_WithPrimer(i) = 100*(TotalProduct_WithPrimer(i)-min(TotalProduct_WithPrimer))/(max(TotalProduct_WithPrimer)-min(TotalProduct_WithPrimer));
end
% Plot normalized data

figure;
semilogx(Inhibitor_Conc,TotalNormalized,'k--');
hold on
title('Normalized RT inhibition data')
xlabel('TFV-DP Concentration [M]')
ylabel('Normalized fluorescence (%)')
legend('Theory, (full-length + fragment) cDNA + ssDNA');
%
```
